# Artesunate Alleviate Kidney Fibrosis by Restoring Klotho Protein and Suppressing Wnt/β-Catenin Signalling Pathway

**DOI:** 10.1101/2025.03.20.644310

**Authors:** Goran H Mohammad, Julius Kieswich, Abhishek Kumar, Kieran McCafferty, Ramyangshu Chakraborty, Christoph Thiemermann, Magdi M Yaqoob

## Abstract

Chronic kidney disease (CKD) is a global health concern that often progresses to renal failure and premature death. Regardless of etiology of CKD, kidney fibrosis is the main determinant of progressive CKD. Renal fibrosis is characterized by excessive collagen and extracellular-matrix (ECM) deposition, which impairs renal function with an irreversible loss of nephrons. Currently, there are no effective antifibrotic therapies to halt the progression of CKD to the end-stage kidney failure (ESKF). Artesunate has recently shown antifibrotic effects in various animal models, but its efficacy in renal fibrosis remains unexplored. In this study, the efficacy of artesunate was evaluated in a unilateral ureteral obstruction (UUO) mouse model and in primary human kidney fibroblasts (HKF). Mechanistic investigation including immunoblot analysis, immunohistochemistry, gene expression assay, enzyme-linked immunosorbent assay (ELISA) and other tools were used to study the underlying molecular mechanisms of antifibrotic effects of artesunate. Results of this study showed that artesunate ameliorated multiple profibrotic pathways including transforming growth factor-beta (TGF-β) expression in UUO model and reduced profibrotic markers including alpha-smooth muscle actin (α-SMA), fibronectin, collagen I, and vimentin in both *in-vivo* and *in-vitro* models. Mechanistic studies indicated that artesunate treatment abrogated the TGF-β/SMAD pathway, restored klotho-protein expression and attenuated both PI3K/Akt and Wnt/β-catenin pathways. Additionally, artesunate inhibited cell proliferation in UUO and induced ferroptosis in HKF cell culture. In conclusions our study demonstrates that artesunate treatment abrogated fibroblast activation, attenuated canonical and non-canonical TGF-β pathways, inhibited cell proliferation in UUO and selectively induced ferroptosis in HKF cell culture, which may offer a potential treatment to attenuate kidney fibrosis and progressive CKD.

**Graphic Abstract:** 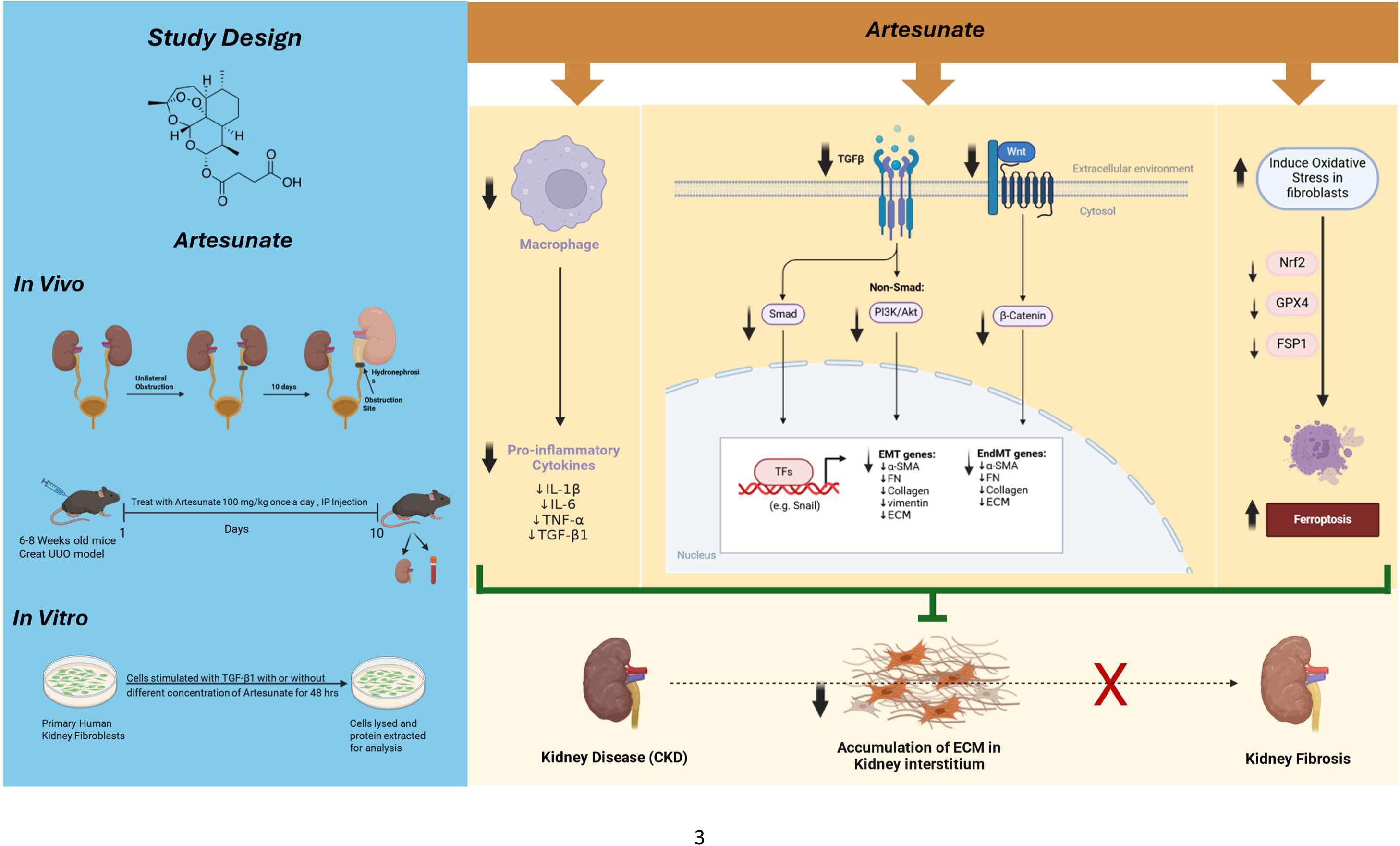

## Introduction

CKD is a global health problem with high co-morbidities and mortality. The disease has a serious risk to human health and often progress to renal failure and premature death^1,2^. Regardless of CKD etiology, kidney fibrosis represents the final pathway of renal demise and is characterised by excessive collagen deposition and ECM accumulation in the tubulointerstitium, causing renal impairment and irreversible nephrons loss^3,4^. The pathogenesis of kidney fibrosis is complex and not fully understood. However, renal injury, inflammation, fibroblast activation and proliferation, growth factors and dysregulation of fibrogenic signalling are thought to be involved^5,6^.

TGF-β and SMAD family signalling are master regulators of renal fibrosis through canonical and non-canonical TGF-β/SMAD pathways^7,8^. In the canonical pathway, TGF-β binds to the TGF-β receptors and triggers SMAD2/3 phosphorylation, then SMAD4 (co-SMAD) directly interacts with phospho-SMAD2/3 and forms a SMAD2/3/4 complex and then translocate to the nucleus to induce and regulate profibrotic genes transcription^7,9^.

The PI3K/Akt/mTOR pathway is a non-canonical TGF-β signalling pathway that contributes to the induction of epithelial-to-mesenchymal-transition (EMT) and renal fibrosis^10,11^. As previously reported, TGF-β activates the PI3K/Akt pathway through direct interaction between the p85-subunit of PI3K and TGF-β receptor II. Upon activation, PI3K phosphorylates its downstream effector, Akt. The PI3K/Akt pathway then activates another downstream effector of the mTOR protein, which is a key regulator of protein synthesis via phosphorylation of S6-kinase (S6K) and eukaryotic initiation factor 4E-binding-protein-1 (4E-BP1). Phosphorylation of S6K and 4E-BP1 by mTOR enhances translational capacity, induces cell proliferation, protein synthesis, EMT induction and renal fibrosis^4,12,13^.

The Wnt/β-catenin pathway is another downstream of TGF-β/SMAD and plays an important role in renal fibrosis and is significantly upregulated during kidney fibrosis^13,14^. During renal fibrosis, the upregulation of Wnt expression activates the Wnt/β-catenin pathway through the binding of Wnt ligands to Frizzled receptors and LRP5/6 co-receptors. This interaction inhibits the β-catenin destruction complex (composed of APC, Axin, GSK-3β, and CK1), leading to the stabilisation and accumulation of β-catenin in the cytoplasm which then facilitates the translocation into the nucleus to interact with TCF/LEF transcription factors that regulate the expression of Wnt-target genes^13–15^. Previous studies have reported the crosstalk between TGF-β/SMAD and Wnt/β-catenin pathways. TGF-β-induced fibrosis can be exacerbated by activation of Wnt/β-catenin pathway, and contributes to the fibroblast activation, ECM and collagen deposition which further aggravates renal fibrosis^16,17^. Additionally, klotho is an anti-aging protein, functioning as an antagonist to the Wnt/β-catenin pathway and often downregulated in CKD. Previous studies have reported the protective role of klotho against kidney injury and fibrosis, revealing that the loss of klotho promotes the Wnt/β-catenin pathway, exacerbates kidney injury and accelerates the progression of CKD to ESRD and kidney fibrosis^18–20^. Indeed, inflammation is another key contributor to the development and progression of renal fibrosis through a complex interplay of cellular and molecular mechanisms. Pro-inflammatory cytokines and growth factors stimulate the fibroblast proliferation and differentiation into myofibroblasts, leading to the excessive ECM production, collagen deposition and tissue scarring^21–23^.

Currently, there are no effective antifibrotic treatments to halt the progression of CKD to the ESKF. Several therapeutic strategies designed to inhibit TGF-β expression, alleviate TGF-β signalling and abrogate the fibroblast activation; however, none of these strategies were translated clinically as demonstrated by unsuccessful clinical trials^6,9,24^. Another potential approach to prevent tissue fibrosis is the inhibition of fibroblast proliferation and transdifferentiation into myofibroblast. Artemisinin is an active ingredient of *Artemisia annua-L*-plant (Chinese herbal medicinal plant), artesunate is one of the Artemisinin derivatives and a first line anti-malaria drug. Artesunate has shown an anti-fibrotic effect in multiple animal models such as ocular^25^, liver^26^ and epidural fibrosis^27^ by inhibiting fibrotic signalling and inducing ferroptosis. However, the efficacy of artesunate in the setting of CKD and renal fibrosis has not been explored and is, therefore, evaluated in this study.

Ferroptosis is a form of regulated cell death characterized by the accumulation of lipid peroxides and iron-dependent oxidative damage. It is driven by the failure of the glutathione dependent antioxidant defence system, particularly the depletion of glutathione and the inactivation of nuclear factor erythroid-2 related factor-2 (Nrf2), glutathione peroxidase-4 (GPX4) and ferroptosis suppressor protein-1 (FSP1). This leads to the unchecked accumulation of reactive oxygen species (ROS) and lipid peroxides, ultimately causing cell death^28,29^.

The aim of the present study is to explore the potential therapeutic effect of artesunate on UUO induced renal fibrosis and it’s potential to prevent the progression of CKD into kidney fibrosis. In addition, we investigate fibrogenic signalling pathways, including TGF-β/SMAD, PI3K/Akt/mTOR and WNT/β-catenin, to elucidate the underlying molecular mechanisms.

## Material and methods

### Animals Experiments

All animal experiments were conducted in accordance with the United Kingdom Home Office Animals 1986 Scientific Procedures with approval granted by our local ethical committee (Project License number: P73DE7999). Male C57BL/6J mice weighing between 20-25g at 6-8 weeks of age were purchased from Charles River UK Ltd. (Margate, Kent, UK) and were used in this study. The mice underwent UUO surgery according to the previous published methods^30,31^. Briefly, mice were anesthetised with isoflurane and the abdominal cavity was exposed using midline laparotomy. Subsequently, the left ureter was isolated and tied off 0.5cm from the pelvis using a sterile 5-0 silk-braided suture, whilst the right ureter was left unclamped. All incisions were closed using a 5-0 Proline suture. The sham operation mice underwent the same surgery but without the ligation of the ureter. Following UUO surgery for a period of 10 days in total, artesunate (100 mg/kg) or vehicle (5% NaHCO3) was administered daily by intraperitoneal injection (100μl/animal). Artesunate (Thermo Scientific, Fisher Scientific, Loughborough, Leicestershire, UK) solution originally was prepared for in vivo injection by dissolving in 5%NaHCO3 (VWR Chemicals BDH, Lutterworth, Leicestershire UK) at a concentration of 100 mg/mL. After surgery, the mice divided into four groups. (1) Sham-treated with vehicle (n=6 mice), (2) Sham-treated with artesunate (n=6 mice), (3) UUO-treated with vehicle (n=8 mice), (4) UUO-treated with artesunate (n=8 mice).

At the end of the experiment, the mice were sacrificed, and both kidneys and blood were collected for further analysis. Both kidneys from each animal were cut in half longitudinally. One half of each kidney was snap-frozen in liquid nitrogen and subsequently stored at -80°C for western blotting analysis. The other half was fixed with 10% Formalin (Merck Life Science UK Ltd, Gillingham, Dorset, UK) for 16h at 4°C then transferred into a 70%v/v ethanol solution for a further 24h and was then subsequently embedded in paraffin for immunohistochemistry. Blood was immediately centrifuged at 10,000 rpm at 4°C for 15 mins and the serum was separated and collected in fresh Eppendorf tubes and stored at -80°C for later analysis.

### Immunohistochemistry

Formalin fixed kidney tissues were embedded in paraffin and cut into 4µm sections. Immunohistochemistry was performed as previously described^32,33^.Briefly, kidney sections were pre-heated in an oven for 1h at 60°C, and then deparaffinised in xylene and hydrated through a series of graded ethanol concentrations (70%-95%). The sections were subjected to heat-mediated antigen retrieval with citrate buffer (pH 4.0) in an autoclave. Endogenous peroxidase activity was inhibited by 3% hydrogen peroxide for 20 mins, followed by incubation with 3% normal horse blocking serum for 20 mins. The sections were then incubated with primary antibody overnight at 4°C for α-SMA, F4/80 and Ki-67 (Antibodies detail listed in Table1). After three washes with PBS containing 0.5% Tween 20, the sections were incubated with horseradish peroxidase (HRP)–conjugated secondary antibody for 30 mins. Staining was visualized using the 3, 3’-diaminobenzidine (DAB) detection system kit. The sections were placed in haematoxylin for 2 minutes, then gently washed and mounted.

Staining with hematoxylin and eosin or sirius red were performed as previously described (35). Images of stained sections were captured using a NanoZoomer S210 Slide Scanner microscope. Ten images of kidney cortex were captured at 20x magnification per mouse and staining was quantified as percentage of total area using ImageJ 1.5t (National Institutes of Health). For Ki67 staining, positive nuclei per field of view were counted instead.

### Western Blot

Immunoblot analysis of kidney samples and human primary kidney cell lines were evaluated by Western Blotting as previously described^34^.Briefly, following protein extraction, protein concentration was measured by the Bicinchoninic Acid (BCA) assay kit (Thermo Scientific, Fisher Scientific, Loughborough, Leicestershire, UK ) and 30 μg protein was run on a precast gel (NuPAGE Novex 4–12% Bis-Tris 1.0 mm, 12 well gel, Invitrogen, Fisher Scientific) and transferred onto a 0.45μm pore size Polyvinylidene fluoride (PVDF) membrane (Amersham, VWR). The membrane was blocked with 5% skim milk solution (OXOID, Merck Life Science Ltd., UK) and incubated with primary antibodies overnight at 4°C (Antibodies detail listed in Supplementary table 1). The membrane was then incubated with appropriate HRP-conjugated secondary antibodies either anti-mouse or anti-rabbit. The antigen antibody reaction was detected by enhanced chemiluminescence substrate (Amersham, VWR). GAPDH or β-actin antibody was used as a protein loading control. Image J software was used to determine relative band intensity in order to quantity protein expression

### Cell cultures

Primary human kidney fibroblasts (HKF) were purchased from DV Biologics and the human renal proximal tubule cells (HK2) were obtained from ATCC. HKF and HK2 cells were grown in DMEM and DMEM-F12 respectively, supplemented with 10% FBS, 1% penicillin/streptomycin (Merck) and the cells were maintained in humidified atmosphere of 20% O2, 5% CO2 at 37 °C as described previously^4,32^. To study the effect of artesunate on HKF and HK2 cell proliferation, the cells were depleted with serum free medium for 24 h, then were stimulated with 10ng/ml TGF-β1 (Rockland, from Cambridge Bioscience, Cambridge UK), before treatment with different concertation of artesunate ranged from 0-120µM. After 48h treatment, the cells washed twice with ice cold PBS and lysed with RIPA buffer (EMD Millipore, Merck) containing protease (Merck) and phosphatase inhibitor cocktails (Roche, Merck). Protein quantity was measured using BCA method and stored at -80°C for further analysis.

### Cell Viability Assay

The cytotoxicity of artesunate treatment on HKF and HK2 cells were assessed by MTS assay (Promega UK Ltd, Southampton, Hampshire UK). The cells were seeded at a density of 5×103 cells per well, in 96 well plates in complete media and placed overnight at 37^°^C in a humidified incubator. The following day, the media was replaced with fresh serum free media containing different concentrations of artesunate ranged from 0-300µM. After 48h incubation, cell viability was measured using MTS assay according to the manufactory instruction. The drugs’ half inhibitory concentration (IC_50_) was calculated by using GraphPad Prism Software (v10) from the dose-response curves of percentage of cell growth vs. drug concentration. Each artesunate treatment concentration was assayed in quadruplicates and the experiment was repeated at least three times.

### ELISA

Pro-inflammatory cytokines in serum samples were evaluated using enzyme linked immunosorbent assay kit (ELISA). The concentration of IL-1β, IL-6 and TNF-α were measured in mice serum, according to the manufacturer’s instruction (IL-1β, Abcam), (IL-6, R&D systems), (TNF-α, Fisher Scientific). The optical density of each cytokine was measured with a microplate reader at 450nm, then the concentration was calculated against a standard curve using GraphPad Prism software (v10).

### Gene Expression Assay

Total RNA was isolated from kidney tissues, and HKF using the RNA extraction kit (Qiagen, Manchester UK) according to the manufacturer’s instruction. One microgram of extracted RNA was reverse transcribed to cDNA using a reverse transcription kit (Qiagen). Real-time PCRs were performed using SYBR Green master mix (BIO-RAD) using 10ng of cDNA. The primers used for the target genes are listed in Supplementary table 2. The relative gene expression was determined by normalising the expression of each gene to that of the β-Actin or GAPDH gene using the 2−ΔΔCt method.

### Statistics

Data are expressed as mean ± SEM. An unpaired student *t-*test was used for comparison between two groups. For multiple comparisons, one- or two-way ANOVA with Tukey’s *post-hoc* test was performed using GraphPad Prism v10. Statistical significance was set at p<0.05.

## Results

### Artesunate reduced pro-fibrotic markers expression, ECM and collagen deposition in both *in-vivo* and *in-vitro* renal fibrosis models

To evaluate the potential therapeutic effect of artesunate, mice were treated with artesunate (100mg/kg/day) for 10 days following UUO surgery. UUO-kidneys exhibited severe dilation, tubulointerstitial expansion and morphological changes. However, kidneys from artesunate-treated mice displayed significantly less dilation and fewer morphological abnormalities (Figure-1A,1E). Additionally, kidney weight index was significantly higher in the UUO groups compared to the sham-groups. However, artesunate treatment significantly reduced the kidney weight index compared with that in the UUO vehicle-treated group (Figure-1E,1F).

**Figure 1.**
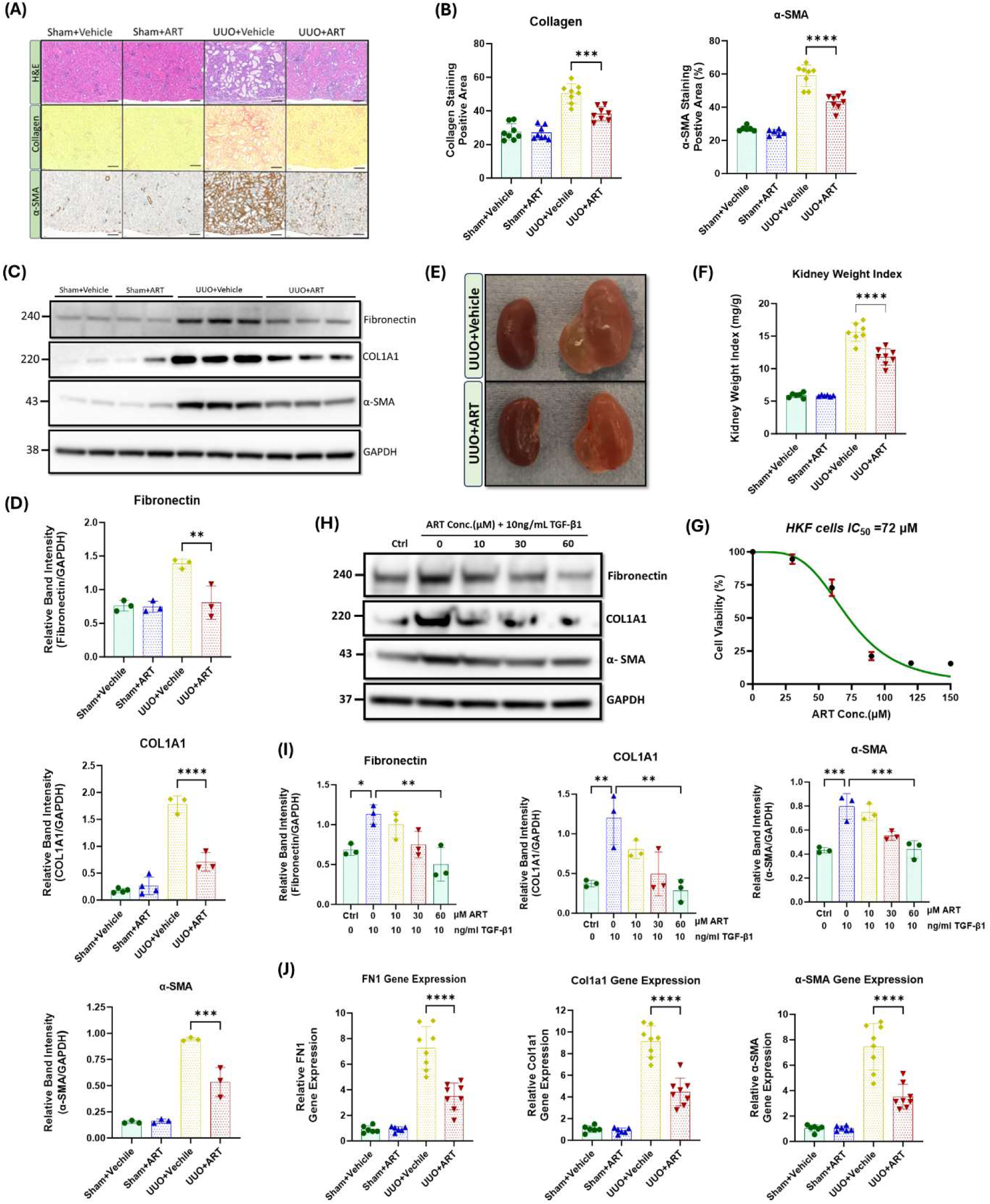
Artesunate treatment ameliorated expression of pro-fibrotic markers and reduced the deposition of ECM and collagen in both *in-vivo* and *in-vitro* renal fibrosis models. **(A)** Staining with Haematoxylin and Eosin (H&E), Sirius Red and immunostaining of kidney sections for α-SMA from obstructed (UUO; 10 days after surgery) or contralateral sham-operated kidneys (control) at 20x magnification (scale bar 50µm). Animals received artesunate (100 mg/kg/day for 10 days) or vehicle as indicated. **(B)** Quantification of collagen deposition and positive α-SMA stained area as percentage of total area. **(C)** Western blot of whole-kidney lysates for fibronectin, collagen and α-SMA expression. PVDF membranes were subsequently stripped of antibodies and reprobed for GAPDH, which served as a protein loading control. **(D)** Relative band intensities of fibronectin, collagen and α-SMA expression were quantified and normalised against GAPDH using ImageJ software. **(E)** Representative images of the obstructed (left) and contralateral kidney (right) in artesunate and vehicle treated group after 10 days post-UUO surgery. **(F)** Effect of artesunate and vehicle treatment on UUO kidney mass was represented by the kidney weight/body weight ratio (kidney weight ratio). Primary human kidney fibroblasts (HKF) were grown in a complete DMEM medium and maintained in a humidified atmosphere incubator. To study the effect of artesunate on HKF cell-proliferation, the cells starved with serum free medium for 24h and then stimulated with 10ng/ml TGF-β1, before treatment with different concertation of artesunate ranged from 0-150mM for 48h. **(G)** The cytotoxicity of artesunate treatment on primary human kidney fibroblast cells was measured by MTS assay and the IC_50_ was calculated. **(H) and (I)** The expression level of fibronectin, collagen and α-SMA protein in response to different artesunate concentrations on HKF cells were assessed using western blotting analysis and the blots were quantified by using ImageJ software and normalised to the GAPDH. **(J)** Representative of relative fibronectin, collagen and α-SMA gene expression from kidney of animals subjected to UUO, treated with artesunate and vehicle as indicated verses the control group. For all graphs, error bars represent the means ± SEM of data from 3–8 animals per group. Student *t*-test or one-way ANOVA was used for statistical analysis. *P<0.05, **p<0.01, ***p<0.001.

Upregulation of profibrotic markers such as collagen, α-SMA and fibronectin are a hallmark of kidney fibrosis. UUO-kidney sections staining showed that the artesunate-treated mice exhibited significantly less collagen deposition in the kidney interstitium and positive α-SMA staining was significantly reduced compared to the UUO-kidneys in vehicle-treated mice (Figure-1A,1B). Western blotting analysis confirmed the effect of artesunate upon profibrotic markers expression which showed reductions of collagen deposition, α-SMA and fibronectin expression in UUO-kidneys by 60%, 43% and 42% respectively, compared to vehicle-treated UUO-kidneys (Figure-1C,1D). Our *in-vitro* results further confirmed the therapeutic effect of artesunate with significantly reduced profibrotic markers present in renal fibroblast cell culture. HKF cells were stimulated with TGF-β in the presence or absence of artesunate. First, a dose response curve of artesunate was performed by MTS assay and IC_50_ was calculated as 72μM (Figure-1G). Lower doses than the IC_50_ have been chosen for this study. Western blotting analysis revealed that artesunate reduced profibrotic markers expression in a dose-dependent manner in renal fibroblasts. At a concentration of 60μM, artesunate significantly downregulated fibronectin, collagen, α-SMA, connective tissue growth factor (CTGF) and vimentin expression by 55%, 76%, 45%, 44% and 47% respectively and inhibited renal fibroblast proliferation (Figure-1H,1I and Suppl.Fig.1)

Gene expression assay further confirmed the ability of artesunate to prevent the progression of kidney fibrosis. Artesunate treatment significantly reduced fibronectin, collagen, α-SMA, CTGF and vimentin gene expression in UUO-kidneys by 52%, 51%, 53%, 51% and 52% respectively (Figure-1J and Suppl.Fig.2).

**Figure 2.**
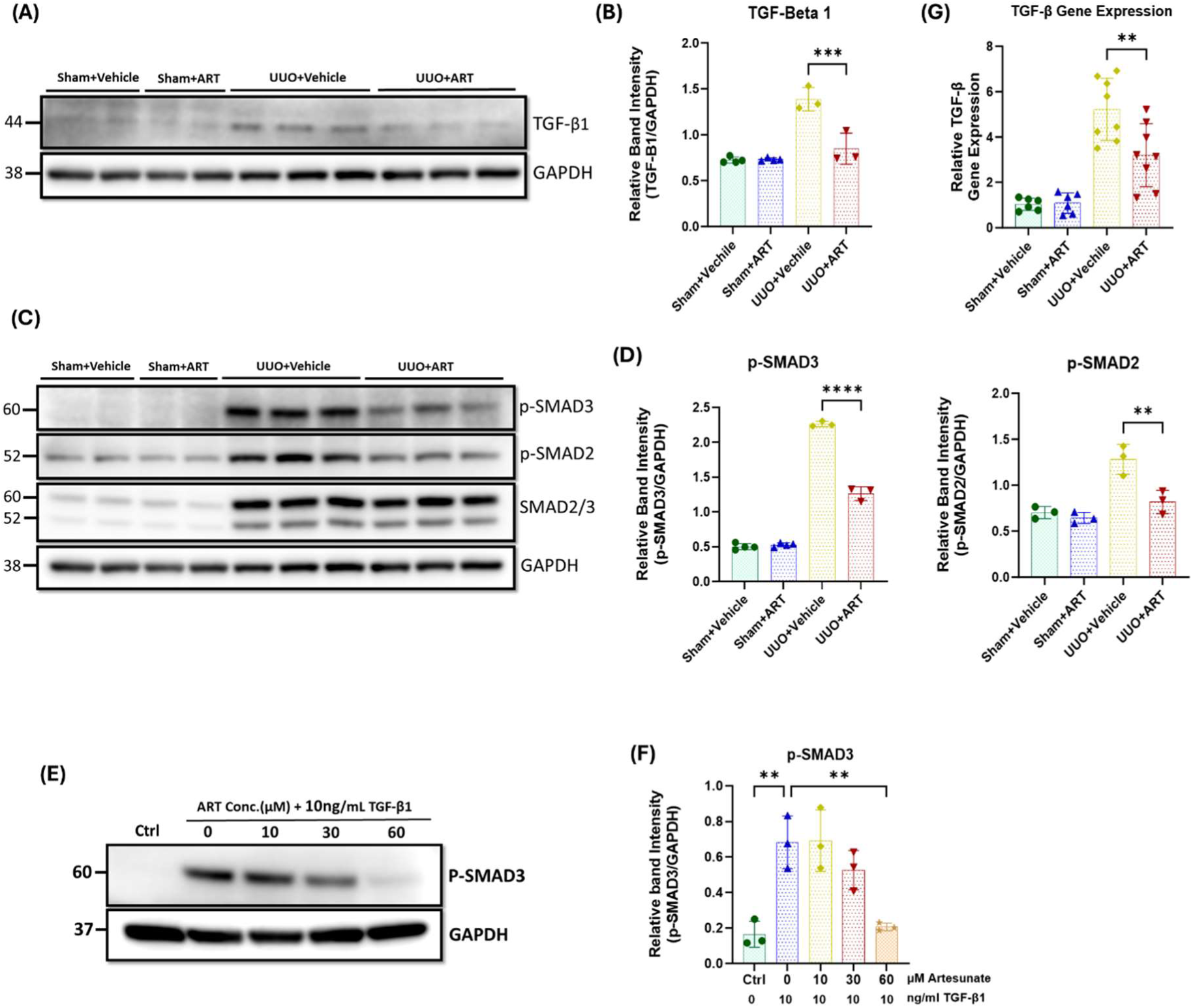
Artesunate administration reduced TGF-β expression and attenuated TGF-β/SMAD pathway in UUO kidneys and renal fibroblast cell culture. (A) and. **(B)** the expression level of TGF-β protein in UUO kidneys and control group were assessed using western blotting technique and the blots were quantified by using ImageJ software and normalised to GAPDH. **(C) and (D**) Western blotting was performed to evaluate the expression level of p-SMAD3, p-SMAD2 and SMAD2/3 in UUO kidneys and control group and ImageJ software was used to quantify the blots and normalised to GAPDH. **(E) and (F)** HKF cells were starved with serum free medium for 24h and then stimulated with 10ng/ml TGF-β1, before treatment with different concertation of artesunate ranged from 0-60mM for 48h. The cells were then lysed and the expression level of p-SMAD3 protein was determined by Western blotting, the blots were quantified by using ImageJ software and normalised to the GAPDH. **(G)** mRNA level of TGF-β in UUO kidneys and control group were quantified by using quantitative polymerase chain reaction (qPCR). The data are presented as the means ± SEM from 3–8 animals per group. Student *t*-test or one-way ANOVA was used for statistical analysis. *P<0.05, **p<0.01, ***p<0.001.

Overall, these results demonstrate the ability of artesunate to supress the expression of pro-fibrotic markers and reduce ECM and collagen deposition in both *in-vivo* and *in-vitro* renal fibrosis models and to inhibit renal fibroblasts expansion and transdifferentiation into myofibroblasts.

### Artesunate suppressed TGF-β expression and alleviated the activation of the TGF-β/SMAD pathway in UUO-kidneys and renal fibroblasts

TGF-β plays a central role in the development and progression of kidney fibrosis through canonical and non-canonical pathways. Consequently, we investigated the effect of artesunate treatment on TGF-β expression and the TGF-β/SMAD pathway.

Western blotting analysis revealed a dramatic upregulation of TGF-β expression and increased phosphorylation of both SMAD2 and SMAD3 in obstructed kidneys. However, artesunate administration effectively suppressed TGF-β expression and significantly reduced the SMAD2 and SMAD3 phosphorylation by 39%, 32% and 44%, respectively (Figure-2A-2D).

SMAD3 is a key downstream mediator of the TGF-β signalling, and a critical driver of the fibrotic process. It facilitates fibroblast differentiation into myofibroblasts and directly promotes the expression of fibrogenic genes^35–38^. Western blotting analysis showed a dramatic upregulation of phospho-SMAD3 in stimulated fibroblast cell culture, whilst artesunate treatment gradually reduced phospho-SMAD3 expression in a dose-dependent manner, restoring it to control levels at a concentration of 60µM (Figure-2E,2F).

Gene expression assay confirmed the capability of artesunate to reduce TGF-β mRNA expression in UUO-kidneys. As shown in the figure 2G, artesunate administration remarkably reduced TGF-β mRNA-expression by 39% in UUO-kidneys compared to the vehicle-treated group.

Collectively, these results demonstrate the ability of artesunate to supress TGF-β expression and alleviate the activation of the TGF-β/SMAD signalling and repress the transcription of pro-fibrotic genes in UUO-kidneys and renal fibroblast cell culture.

### Artesunate attenuated the PI3K/Akt pathway and reduced cell proliferation in UUO-kidneys and renal fibroblasts

To understand the molecular mechanisms underlying the inhibitory effect of artesunate in obstructive kidney and renal fibroblasts, we next investigated the PI3K/Akt pathway. As shown in figure-3A, phospho-Akt expression dramatically increased in UUO-kidneys, whilst artesunate administration significantly reduced the phosphorylation of Akt by 52%.

**Figure 3.**
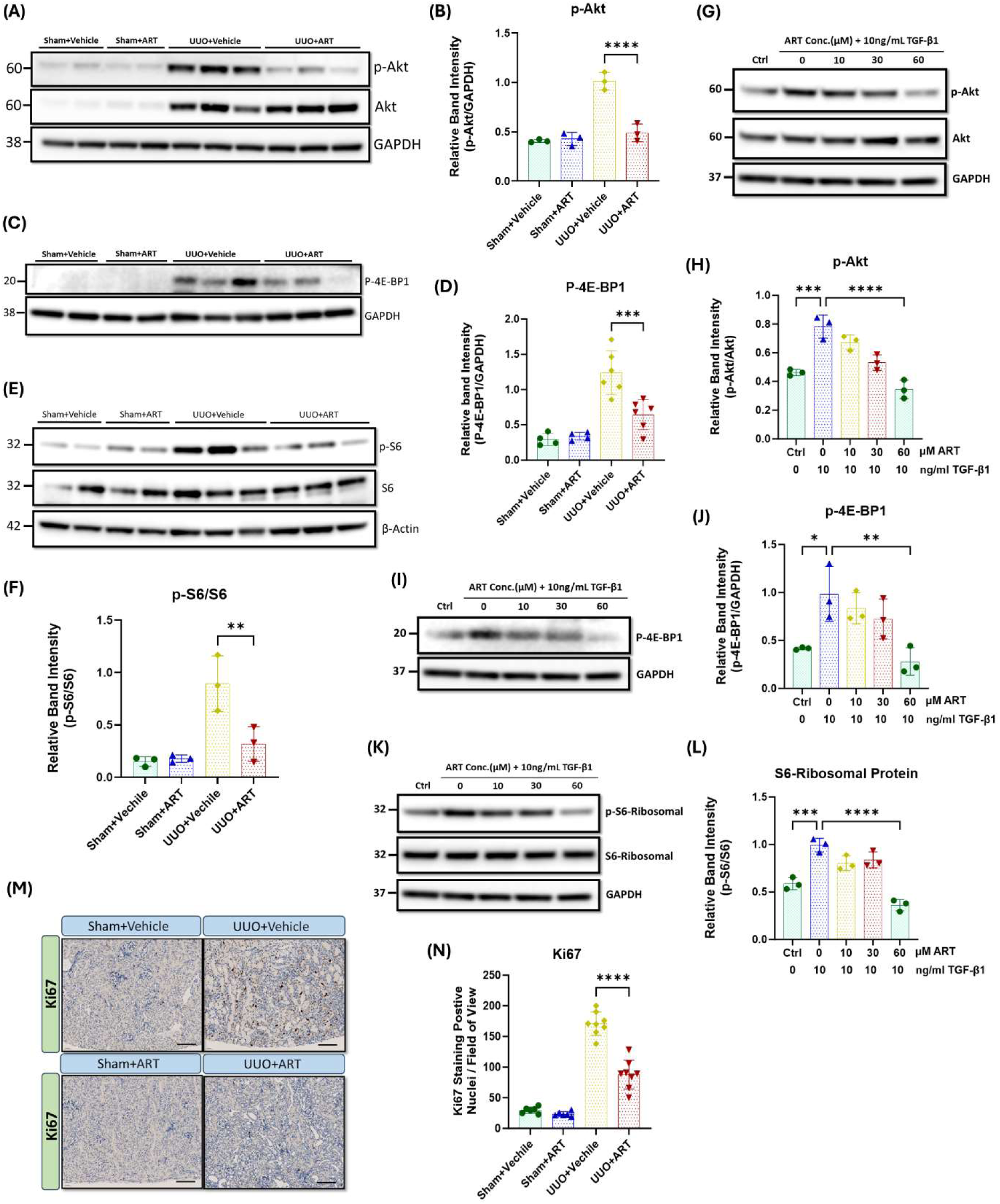
Artesunate administration ameliorated PI3K/Akt pathway and reduced cell-proliferation in UUO kidneys and renal fibroblast cell culture. (A) and. **(B**) p-AKT and AKT expression were evaluated in UUO kidneys and control group by using western blotting analysis and ImageJ software was used to quantify the blots and normalised to GAPDH protein expression **(C) and (D)** Expression of p-4E-BP1 protein was assessed in UUO kidneys and control group by western blotting and ImageJ software was used to quantify the blots normalised to GAPDH. **(E) and (F)** Western blotting was performed to assess the expression level of p-S6K and S6K protein in UUO kidneys and control group and ImageJ software was used to quantify the blots and normalised to β-Actin. HKF cells were starved with serum free medium for 24h and then stimulated with 10ng/ml TGF-β1, before treatment with different concertation of artesunate ranged from 0-60mM for 48h. The cells were then lysed and the expression level of p-Akt, Akt **(G,H)**, p-4E-BP1 **(I,J)**, p-S6k and S6K **(K,L)** proteins were determined by western blot. ImageJ software was used to quantify the blots and p-Akt and p-S6 normalised to the total Akt and S6K protein respectively and p-4E-BP1 normalised to GAPDH. **(M) and (N)** Representative UUO kidney and control kidney sections were immunostained with anti-Ki67 antibody and the number of Ki67 positive nuclei per field of view in UUO kidney and control kidney sections were quantified (20x magnification, scale bar 50µm). The data are presented as the means ± SEM from 3–8 animals per group. Student *t*-test or one-way ANOVA was used for statistical analysis. *P<0.05, **p<0.01, ***p<0.001.

Akt phosphorylation is a critical step in the activation of several downstream signalling pathways including mTORC1 pathway and promotes cell proliferation, migration and survival by phosphorylating key downstream targets, including S6K and 4E-BP1 ^12,13^. Western blotting analysis showed a dramatic upregulation of phospho-S6K and phospho-4E-BP1 in UUO-kidneys, whilst artesunate treatment significantly reduced S6K and 4E-BP1 phosphorylation by 64% and 48% respectively (Figure-3C-3F). Consistent with *in-vivo* findings, our *in-vitro* results showed a reduction of Akt phosphorylation in a dose-dependent manner in response to the artesunate treatment as demonstrated by western blot. At a concentration of 60μM for 48h, artesunate significantly reduced phosphorylation of Akt, S6K and 4E-BP1 by 56%, 64% and 72% respectively, compared to the untreated fibroblasts, aligning with the expressions shown in the control group (Figure-3G-3L).

Moreover, inhibition of fibroblasts proliferation was further confirmed by immunostaining of kidney sections with Ki67, a proliferation marker. Ki67-positive nuclei were scarcely detected in the sham-kidneys (Figure-2M). UUO resulted in a dramatic increase in the detection of Ki67-positive nuclei, primarily in the interstitium, whilst artesunate administration significantly reduced the number of proliferating cells by 50% (Figure-2M,2N).

Together, these results demonstrate the ability of artesunate to downregulate the PI3K/Akt/mTORC1 pathway and supress fibroblast expansion and myofibroblast proliferation in UUO-kidneys and renal fibroblasts.

### Artesunate restored klotho expression and attenuated Wnt/β-catenin pathway in UUO-kidneys

To further investigate the inhibitory mechanisms of artesunate on kidney fibrosis in the UUO model, the Wnt/β-catenin pathway was studied. This pathway contributes to the pathogenesis of kidney fibrosis by inducing pro-fibrotic genes, EMT, ECM production and collagen deposition^14,15^.

Glycogen synthase kinase-3-beta (GSK3β) is a key regulator of the Wnt/β-catenin pathway. In the presence of Wnt protein, Wnt-ligand binds to their receptor, leading to the inhibition of GSK3β activity. Inactivation of GSK3β prevents the phosphorylation and subsequent degradation of β-catenin, leading to the cytoplasmic accumulation of β-catenin which then translocation into the nucleus to induce pro-fibrotic gene expression including that for Snail^14,15^. Western blotting analysis showed that the artesunate administration significantly reduces Wnt-1 expression, increases GSK3β phosphorylation, reduces both β-catenin and active β-catenin and Snail-1 expression by 33%, 52%, 59%, 47%, and 36% respectively, in UUO-kidneys compared with that shown in the vehicle-treated group (Figure-4E-4H).

**Figure 4.**
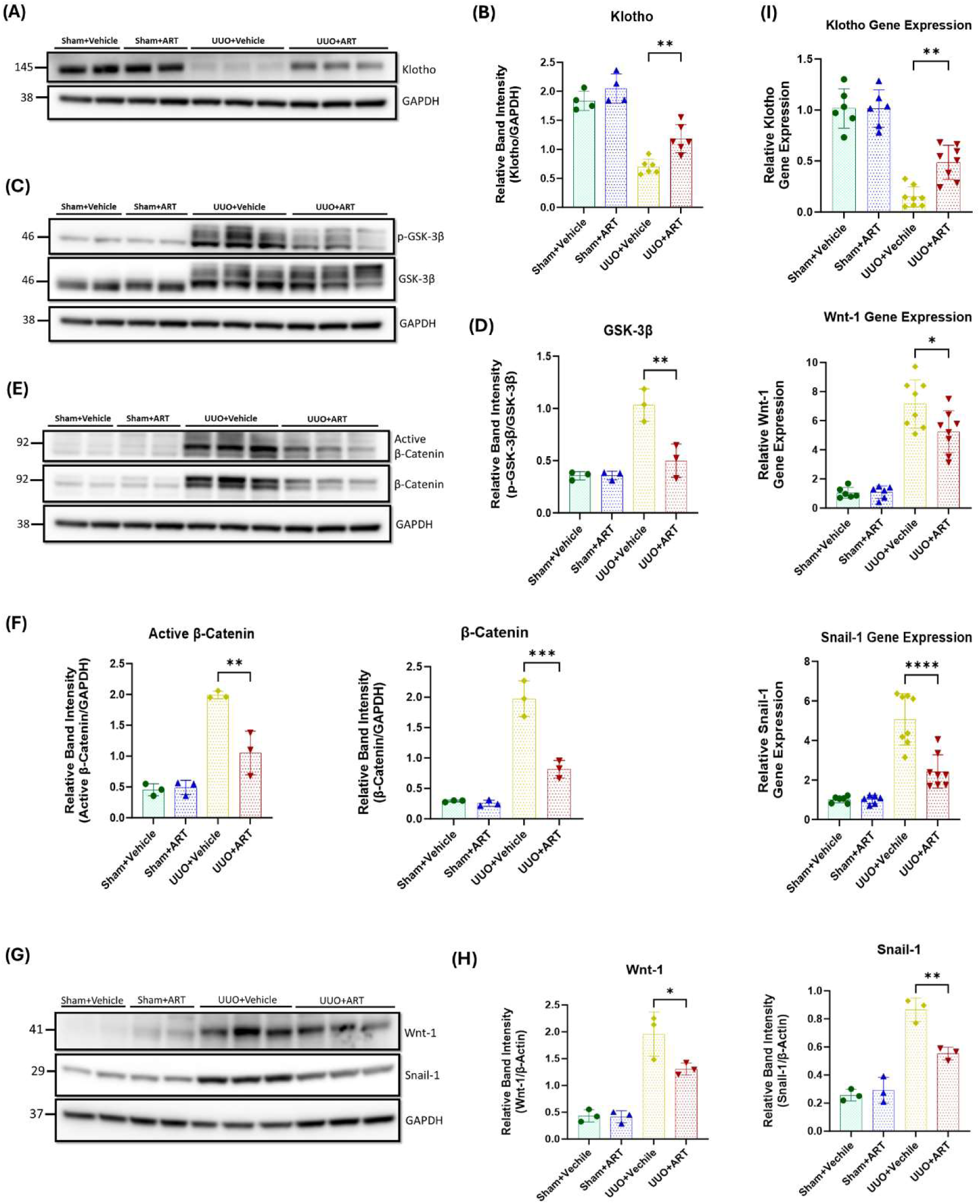
Artesunate administration restored klotho protein and attenuated Wnt/β-Catenin pathway and reduced transcription of fibrosis-related genes in UUO kidney. (A-. **H)** Western blotting was performed to evaluate the expression of Klotho **(A,B)**, p-GSK-3β, GSK-3β **(C,D)**, Active β-Catenin, β-Catenin **(E,F)**, Wnt-1 and Snail-1 **(G,H)** proteins in UUO kidneys and control group. ImageJ software was used to quantify the blots and p-GSK-3β protein expression normalised to the total GSK-3β protein whilst the other proteins expression was normalised to GAPDH. **(I)** mRNA level of klotho, Wnt-1 and Snail-1 in UUO kidneys and sham group were quantified by using qPCR. The data are presented as the means ± SEM from 3–8 animals per group. Student *t*-test or one-way ANOVA was used for statistical analysis. *P<0.05, **p<0.01, ***p<0.001.

Next, we investigated the effect of artesunate on klotho expression. Klotho acts as an antagonist of the Wnt/β-catenin pathway by directly interacting with and sequestering Wnt-ligands, thereby preventing their activation of the receptor complex^18–20^. This inhibition protects the kidney against fibrosis and helps maintain kidney function. As shown in figures 4A and 4B, artesunate treatment not only reduced Wnt-1 protein expression but also partially restored klotho expression in the UUO-kidneys, whilst klotho expression was remarkably reduced in the vehicle treated UUO-kidneys.

Furthermore, gene expression assay confirmed the capability of artesunate to attenuate kidney fibrosis. Artesunate treatment significantly increased klotho mRNA and significantly reduced Wnt-1 and Snail-1 mRNA-expression in UUO-kidneys compared with that shown in vehicle-treated UUO-kidneys (Figure-4I).

Together, these results demonstrate the ability of artesunate to downregulate the Wnt/β-catenin pathway and to restore klotho expression and thereby supressing kidney fibrosis and improve kidney function in the UUO mice model.

### Artesunate alleviated macrophages and pro-inflammatory cytokines in UUO-kidneys

Inflammation is a key driver of fibrotic processes that lead to the development of CKD and ultimately, kidney fibrosis and failure^21–23^. UUO resulted in a dramatic increase in macrophage marker expression, whilst artesunate administration significantly reduced as evaluated by western blot (Figure-5A,5B). Immunostaining of kidney tissues with F40/80, a unique marker of murine macrophages, showed similar results. Artesunate significantly reduced F40/80 expression in UUO-kidneys compared to the UUO-kidneys from the vehicle-treated group (Figure-5C,5D).

**Figure 5.**
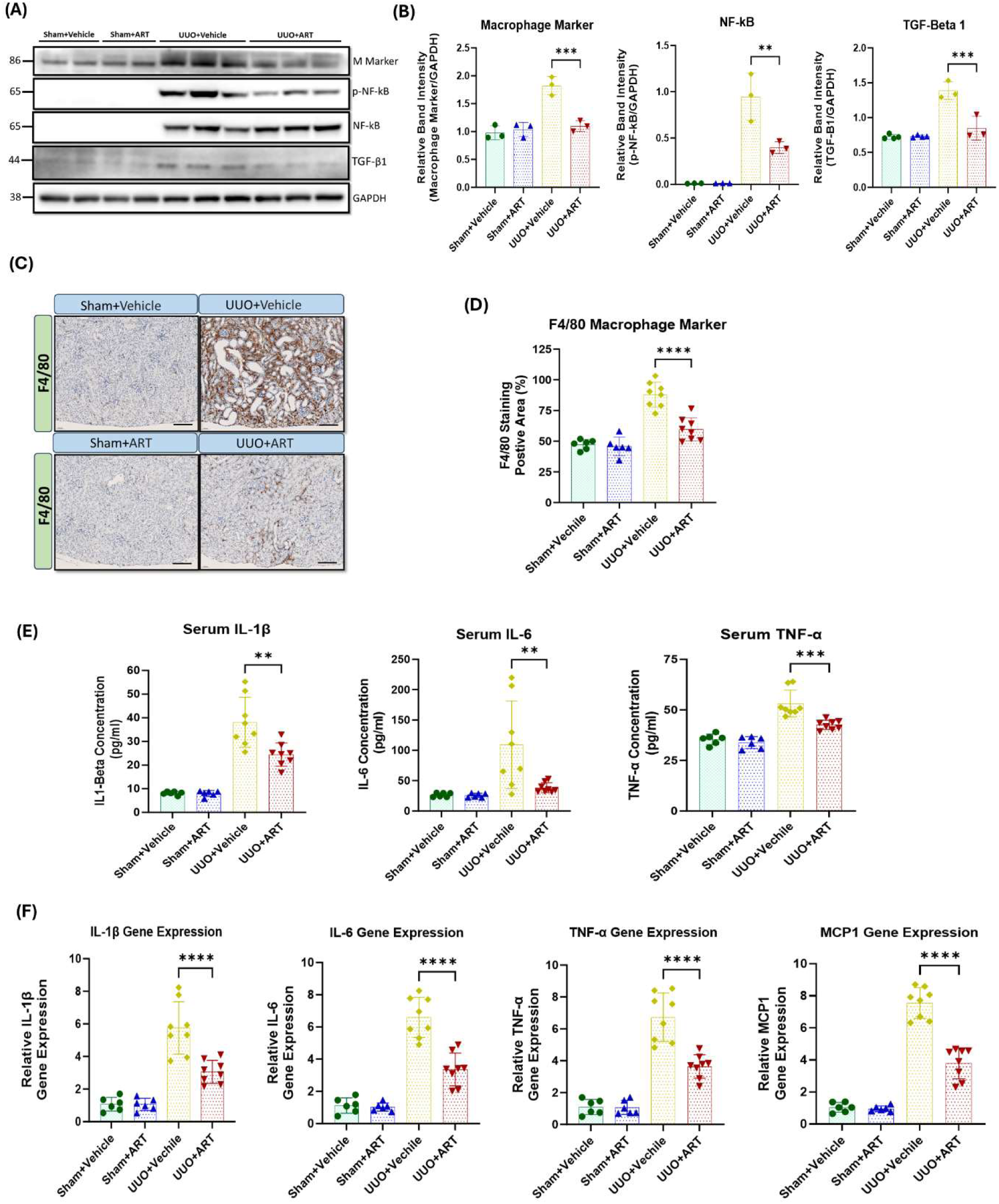
Artesunate administration suppressed macrophage marker and pro-inflammatory cytokine expression in UUO kidney. (A) and. **(B)** the expression level of macrophage marker, p-NF-kB, NF-kB and TGF-β protein in UUO kidneys and control group were evaluated using western blotting technique and the blots were quantified buy using ImageJ software and normalised to the GAPDH. **(C) and (D)** Immunostaining of UUO and control kidney sections for macrophage infiltration with F4/F80 at 20x magnification (scale bar 50µm) and quantification of positively stained area as percentage of total area. **(E)** The serum level of IL-1β, IL-6 and TNF-α cytokines in animals subjected to UUO surgery and sham group were measured by ELISA technique. **(F)** mRNA level of pro-inflammatory cytokines IL-1β, IL-6, TNF-α and MCP1 in UUO kidneys and sham group were quantified by using qPCR. The data are presented as the means ± SEM from 3–8 animals per group. Student *t*-test or one-way ANOVA was used for statistical analysis. *P<0.05, **p<0.01, ***p<0.001.

Nuclear factor kappa B (NF-κB) plays a crucial role in the development of kidney fibrosis by promoting inflammation, fibroblast activation, EMT, and oxidative stress^21–23^. We therefore investigated the effect of artesunate treatment on NF-κB activation and pro-inflammatory markers expression. NF-kB was not detectable in sham-operated kidneys, whereas considerable upregulation of phospho-NF-kB was observed in UUO-kidneys. Artesunate administration significantly suppressed NF-kB phosphorylation by 58% compared to the UUO-kidneys of the vehicle-treated group (Figure-5A,5B). Similarly, artesunate administration significantly reduced serum concentrations of IL-1β, IL-6 and TNF-α by 38%, 65% and 36% respectively, in UUO mice compared to UUO vehicle-treated mice (Figure-5E).

Additionally, gene expression assay demonstrated the capability of artesunate to reduce inflammation in UUO-kidneys and significantly reduced IL-1β, IL-6, TNF-α and MCP1 mRNA-expression by 47%, 49%, 46% and 50% respectively, compared to the UUO-kidneys of the vehicle-treated group (Figure-5F).

Overall, these results showed that the artesunate not only reduced profibrotic protein expression but was also able to downregulate pro-inflammatory cytokines and improve kidney function.

### Artesunate increased oxidative stress and induced ferroptosis in renal fibroblasts

Previous research shows that artesunate induces oxidative stress, elevates ROS, and triggers ferroptosis through several mechanisms^25,26^. Therefore, we initially investigated NADPH oxidase-4 (NOX4) expression in response to the artesunate treatment as it involves ROS production and the induction of oxidative stress. Artesunate significantly elevated NOX4 expression in renal fibroblasts, at a treatment concentration of 30μM for 48h, artesunate increased NOX4 expression by 40% compared to untreated fibroblasts (Figure-6A,6B).

**Figure 6.**
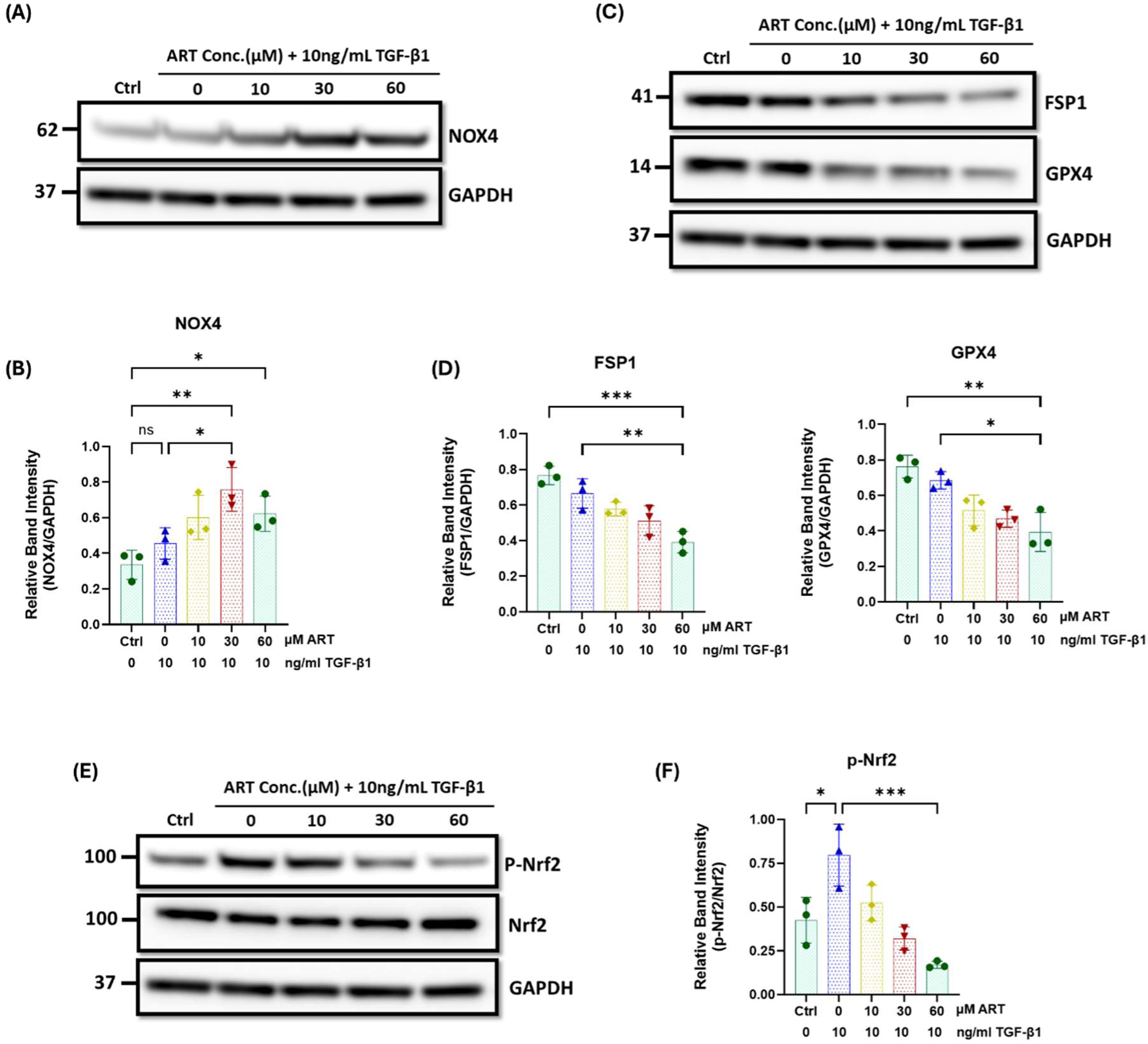
Artesunate treatment increased oxidative stress and induced ferroptosis in human primary kidney fibroblast cells. Serum starved primary human renal fibroblasts were stimulated with 10ng/ml TGF-β1 before treatment with different concertation of artesunate ranged from 0-60μM for 48h. The cells were then lysed with RIPA buffer containing protease and phosphatase inhibitor cocktails. Protein quantity was measured using BCA method and equal protein quantity was used for western blotting analysis. The expression level of NOX4 **(A,B)**, FPS1 and GPX4 **(C,D),** and p-Nrf2 and Nrf2 **(E,F)** in respond to the different concentrations of artesunate were evaluated using western blotting technique and the blots were quantified buy using ImageJ software and p-Nrf2 normalised to the total Nrf2 protein, and all other proteins normalised to the GAPDH. Bars represent the means ± SEM from three independent experiments. Student *t*-test or one-way ANOVA was used for statistical analysis. *P<0.05, **p<0.01, ***p<0.001.

Nrf2 is another important transcription factor that plays a key role in cellular defence against oxidative stress, maintaining redox homeostasis and enhancing the activity of GPX4 and SCL7A11 to protect cells from oxidative damage^39–42^. Western blotting analysis showed that the artesunate treatment gradually reduced Nrf2 phosphorylation and downregulated the expression of FSP1 and GPX4 in a dose-dependent manner in renal fibroblasts (Figure-6C-6F). At a treatment concentration of 60µM for 48h, artesunate significantly reduced Nrf2 phosphorylation, FSP1 and GPX4 expression by 79%, 41% and 42% respectively, compared to the untreated fibroblasts.

In contrast, artesunate treatment did not significantly elevate NOX4 expression, Nrf2 phosphorylation and the expression of GPX4 and FSP1 in human renal proximal tubule cells (Suppl.Fig.3). This finding indicates that artesunate did not increase oxidative stress or induce ferroptosis in HK2 cells.

Overall, these findings suggest that artesunate selectively increases oxidative stress and induces ferroptosis in fibroblasts without affecting other kidney cells. This selective mechanism effectively inhibits fibroblasts proliferation and their transdifferentiation into myofibroblasts, a key mechanism in the pathogenesis of kidney fibrosis.

## Discussion

The most important finding of this study is that artesunate treatment significantly attenuated kidney fibrosis in a UUO mouse model and induced ferroptosis in a renal fibroblast culture. Kidney fibrosis is characterised by excessive accumulation of ECM and collagen in the kidney interstitium. The overproduction of ECM disrupts the normal tissue architecture and impairs kidney function and accelerates the progression to kidney failure. The pathogenesis of kidney fibrosis is very complex and several profibrotic, proinflammatory growth factors and pathways are involved in the progression of this disease^3–6^.

TGF-β plays a pivotal role in the pathogenesis of kidney fibrosis and is consistently upregulated in CKD^7,8^. Consequently, targeting the TGF-β pathway represents a potential therapeutic approach to mitigate the progression of CKD to fibrosis. The therapeutic potential of artesunate and its underlying mechanisms have been investigated in this study. Our findings have demonstrated that artesunate attenuates multiple fibrogenic signalling pathways in both *in-vitro* and *in-vivo* models.

Artesunate has been shown to mitigate TGF-β expression by both canonical and non-canonical TGF-β/SMAD pathways^25^. Chen *et al*. reported that artesunate reduced the expression of TGF-β, SMAD3, and collagen in the type-1 diabetic rat, ultimately attenuating the TGF-β/SMAD pathway and delaying the progression of kidney fibrosis^43^. Similarly, Jingyuan *et al*. demonstrated that artesunate inhibited fibroblast activation by reducing TGF-β expression, decreasing SMAD2/3 phosphorylation and suppressing the TGF-β/SMAD pathway in primary human ocular fibroblasts^25^. Furthermore, Jing *et al*. found that artesunate reduced profibrotic proteins expression including fibronectin, collagen, and α-SMA, thereby mitigating kidney fibrosis in the UUO rat model^44^.

Our findings align with these observations, showing that artesunate administration in UUO mice significantly mitigated TGF-β expression and markedly attenuated the TGF-β/SMAD pathway by reducing SMAD2/3 phosphorylation. Artesunate also alleviated the expression of key fibrotic markers, including fibronectin, collagen, α-SMA and CTGF in both UUO-kidneys and renal fibroblast cell culture and effectively inhibited fibroblast proliferation and differentiation into myofibroblasts.

Additionally, SMAD3 is a key mediator of the TGF-β/SMAD pathway and plays a crucial role in promoting kidney fibrosis^35,37^. As previously reported, pharmacological inhibition of SMAD3 phosphorylation in experimental kidney injury models disrupts the TGF-β/SMAD pathway and significantly reduces the fibrotic response and kidney fibrosis^45,46^. Moreover, genetic ablation of SMAD3 in murine models significantly attenuates renal fibrosis^35–37^. Supporting these observations, our *in-vitro* study reveals SMAD3 as an essential regulator of fibroblast proliferation and myofibroblast differentiation. We observed that artesunate significantly reduced SMAD3 phosphorylation in HKF cells, disrupting the TGF-β/SMAD pathway and potentially mitigating renal fibrosis.

Furthermore, the PI3K/Akt pathway is a downstream target of TGF-β signalling that significantly contributes to renal fibrosis. Our results demonstrated that artesunate administration ameliorated the activation of the PI3K/Akt pathway and significantly reduced Akt phosphorylation in UUO-kidneys. Activated Akt promotes renal fibrosis through activation of mTORC1 and inhibition of GSK-3β^13,47^. mTORC1 is a key downstream effector of AKT; when activated, AKT inhibits tuberous sclerosis complex-2 (TSC2), a negative regulator of mTORC1, resulting in mTORC1 phosphorylation. Consequently, active mTORC1 enhances the translation of fibrosis-related genes by phosphorylating downstream targets like S6K and 4E-BP1^10–13^. Our results showed that artesunate effectively alleviated PI3K/AKT/mTORC1 pathway and inhibited the fibroblast proliferation and differentiation by significantly reduction of Akt, S6K and 4E-BP1 phosphorylation in both UUO-kidneys and fibroblast cell cultures.

Moreover, the Wnt/β-catenin pathway is upregulated in kidney fibrosis and exhibits crosstalk with TGF-β signalling. TGF-β overexpression enhances the expression of Wnt-ligands and β-catenin pathway components and amplifying the fibrotic response^16,17^. We observed a significant upregulation of the Wnt/β-catenin pathway in UUO-kidneys, which was notably suppressed by artesunate administration. Our findings elucidate the mechanisms by which artesunate attenuates Wnt/β-catenin signalling in the UUO model, and we propose four potential mechanisms. Firstly, artesunate significantly reduced TGF-β expression and signalling pathway, subsequently downregulating the Wnt/β-catenin pathway. Secondly, artesunate significantly reduced Wnt expression in the UUO-kidneys, thereby attenuating the Wnt/β-catenin pathway. Thirdly, klotho acts as an antagonist of the Wnt/β-catenin pathway and is downregulated during CKD. Previous studies have reported the protective role of klotho against kidney injury and fibrosis, demonstrating that the loss of klotho exacerbates kidney injury and accelerates the progression of CKD to kidney fibrosis^18,19^. Lili *et al*. reported that the loss of klotho contributes to kidney injury by derepression of the Wnt/β-catenin pathway^18^. Additionally, Qian *et al*. found a klotho-derived peptide (KP1) that protects the kidneys by targeting TGF-β signalling^19^. Furthermore, Wei *et al*. demonstrated that dihydroartemisinin, an artemisinin derivative, suppresses renal fibrosis in a mouse model by inhibiting DNA-methyltransferase activity and enhancing klotho expression^20^. We observed a significant reduction in klotho expression in UUO-kidneys. Notably, artesunate administration not only reversed this effect but also partially restored klotho in the UUO-kidneys. This restoration was associated with the inhibition of Wnt-receptor interaction, thereby attenuating the Wnt/β-catenin pathway. Finally, previous studies have demonstrated crosstalk between the PI3K/Akt and Wnt/β-catenin pathways, with GSK3β serving as a key interaction point^48^. In the absence of Wnt signalling, GSK3β phosphorylates β-catenin at multiple sites, typically Ser33, Ser37, and Thr41, marking it for ubiquitination and subsequent degradation. Conversely, in the presence of activated Akt, it phosphorylates GSK3β at Ser9, thereby inhibiting its activity. Consequently, the PI3K/Akt pathway indirectly activates the Wnt/β-catenin pathway, resulting in accumulation and nuclear translocation of β-catenin, which promotes profibrotic-gene expression^47–51^. Concordant with earlier findings, we observed a significant upregulation of p-Akt, p-GSK3β, and β-catenin in UUO-kidneys whilst artesunate administration reversed these effects.

Inflammation is a key feature of CKD and significantly contributes to disease progression, kidney damage, fibrosis, and systemic complications. Targeting persistent inflammation shows promise in improving CKD patient outcomes^22,23,52,53^. Previous studies have reported the anti-inflammatory effects of artesunate in various models, including UUO^21,43,54^. In line with previous findings, we demonstrated that artesunate significantly reduced pro-inflammatory cytokines IL-1β, IL-6, and TNF-α concentrations, while dramatically downregulated the NF-kB signalling in UUO-kidneys.

Finally, inducing cell death selectively in fibroblasts is a promising therapeutic strategy for treating fibrosis. Previous research demonstrated that treatment with ferroptosis inducers, such as erastin^55,56^ and artemether^57^ alleviates fibrosis progression in liver fibrosis mouse models by triggering ferroptosis in hepatic stellate cells. Kong *et al*. reported that artesunate ameliorates CCl4-induced liver fibrosis by promoting ferroptosis in activated hepatic stellate cells^58^. Likewise, Jingyuan *et al*. found that artesunate treatment protects against ocular fibrosis by suppressing fibroblast activation and inducing mitochondria-dependent ferroptosis^25^. Consistent with previous report, our findings demonstrated that artesunate induces ferroptosis in renal fibroblasts, inhibiting both proliferation and differentiation into myofibroblasts.

The mechanism underlying the induction of ferroptosis by artesunate in fibroblasts remains unclear. However, we identified key mechanisms through which artesunate initiates and activates ferroptosis. Artesunate treatment significantly upregulated NOX4 expression, leading to an increase in oxidative stress and an elevation in ROS production. Consequently, this led to the promotion of lipid peroxidation and the depletion of key antioxidant defences, including GSH and GPX4, which normally protect cells from oxidative damage.

Moreover, artesunate disrupts another antioxidant defence system by attenuating Nrf2 signalling and reducing FSP1 expression in renal fibroblasts. These findings suggest that artesunate inhibits fibroblast proliferation through the induction of ferroptosis by impairing the cellular antioxidant defence mechanisms. In contrast, the same concentration and duration of artesunate treatment did not alter NOX4 expression, Nrf2 phosphorylation, or the expression of FSP1 and GPX4 proteins in HK2 cells. These results suggest that artesunate selectively inhibits fibroblast proliferation and induces ferroptosis specifically in renal fibroblasts, without affecting the viability or function of other kidney cells.

In conclusion, the results of this study demonstrated the potential therapeutic effect of artesunate and the ability to protect the kidney against fibrosis in a renal fibrosis mouse model by abrogation of fibroblast activation, blunting of multiple fibrogenic pathways, and inhibition of cell proliferation. Additionally, in cultured human primary kidney fibroblasts it induced ferroptosis, and caused the inhibition of fibroblasts proliferation and differentiation. Thus, artesunate may offer a potential treatment to attenuate kidney fibrosis and progressive CKD in the future.

## Statements & Declarations

## Acknowledgements

This research was supported by Diabetic Kidney Disease Centre, Renal Unit, Barts Health National Health Service Trust, The Royal London Hospital, London, UK

## Authors Contributions

GM and MMY conceived and designed the study. GM, MMY, SH, KM, CT, JK, RC and AK contributed to the acquisition and/or analysis of the data. GM drafted the manuscript, and all authors reviewed and approved the final version.

## Ethics approval

All animal experiments were conducted in accordance with the United Kingdom Home Office Animals 1986 Scientific Procedures with approval granted by our local ethical committee (Project License number: P73DE7999).

## Consent to participate

Informed consent was obtained from all individual participants included in the study

## Supplemental information

Document S1: Supplementary Tables, Table 1 and 2. Document S2: Supplementary Figures, Figures S1–S3.

## Data and materials availability

The data supporting the findings of this study are available from the corresponding author upon request.

