## Supplementary table for "Artesunate Alleviate Kidney Fibrosis by Restoring Klotho Protein and Suppressing Wnt/β-Catenin Signalling Pathway"

**Supplementary Tables**

**Supplementary table 1:** The list of the Antibodies used in immunohistochemistry and western blotting experiment.

| **Antibody Name** | **Dilution** | **Source** | **Catalogue Number** | **Techniques** | **Manufacture** |
| --- | --- | --- | --- | --- | --- |
| α-SMA | 1:500 | Mouse | A5228 | IHC | Sigma |
| F4/80 | 1:300 | Rabbit | 70076 | IHC | Cell Signaling |
| Ki-67 | 1:300 | Rabbit | ab15580 | IHC | Abcam |
| Fibronectin | 1:1000 | Rabbit | ab2413 | WB | Abcam |
| Collagen I | 1:500 | Rabbit | 91144 | WB | Cell Signaling |
| α-SMA | 1:1000 | Mouse | A5228 | WB | Sigma |
| TGF-β1 | 1:1000 | Rabbit | 3711S | WB | Cell Signaling |
| p-SMAD2 | 1:1000 | Rabbit | 18338T | WB | Cell Signaling |
| p-SMAD3 | 1:1000 | Rabbit | 9520 | WB | Cell Signaling |
| SMAD2/3 | 1:1000 | Rabbit | 3102 | WB | Cell Signaling |
| p-Akt | 1:1000 | Rabbit | 4056S | WB | Cell Signaling |
| Akt | 1:1000 | Rabbit | 9272 | WB | Cell Signaling |
| p-S6Ribosomal protein | 1:1000 | Rabbit | 2211S | WB | Cell Signaling |
| S6 Ribosomal protein | 1:1000 | Mouse | 2317S | WB | Cell Signaling |
| p-4E-BP1 | 1:1000 | Rabbit | 2855S | WB | Cell Signaling |
| Klotho | 1:1000 | Rabbit | PA5-21078 | WB | Invitrogen |
| Wnt-1 | 1:1000 | Mouse | MA5-15544 | WB | Invitrogen |
| p-GSK-3β | 1:1000 | Rabbit | 5558 | WB | Cell Signaling |
| GSK-3β | 1:1000 | Rabbit | 12456 | WB | Cell Signaling |
| Active β-Catenin (non-phospho-S37/T41) | 1:1000 | Rabbit | ab246504 | WB | Abcam |
| β-Catenin | 1:1000 | Mouse | 37447 | WB | Cell Signaling |
| Snail-1 | 1:1000 | Rabbit | 3879 | WB | Cell Signaling |
| Macrophage Marker (MAC387) | 1:500 | Mouse | sc-66204 | WB | Santa Cruz |
| p-NF-kB | 1:1000 | Rabbit | 3033S | WB | Cell Signaling |
| NF-kB | 1:1000 | Rabbit | 8242S | WB | Cell Signaling |
| NOX4 | 1:1000 | Rabbit | SY0214 | WB | Invitrogen |
| GPX4 | 1:1000 | Rabbit | ab125066 | WB | Abcam |
| FSP1 | 1:1000 | Mouse | sc-377120 | WB | Santa Cruz |
| p-Nrf2 | 1:1000 | Rabbit | PA5-67520 | WB | ThermoFisher |
| Nrf2 | 1:1000 | Rabbit | PA5-88084 | WB | ThermoFisher |
| GAPDH | 1:1000 | Rabbit | 2118L | WB | Cell Signaling |
| β-Actin | 1:2000 | Mouse | A5316 | WB | Sigma |
| GAPDH | 1:1000 | Rabbit | 2118L | WB | Cell Signaling |
| α-Tubulin | 1:2000 | Mouse | T6074 | WB | Sigma |

**Supplementary table 2:** The list of the primer sequences used in gene expression assay (qPCR) experiment.

| **Primer Name (Mouse)** | **Forward primer Sequence (5'->3')** | **Reverse primer Sequence (5'->3')** |
| --- | --- | --- |
| Fibronectin (FN1) | ATGTGGACCCCTCCTGATAGT | GCCCAGTGATTTCAGCAAAGG |
| Col 1 a1 | TCGTGTAAACTCCCTCCACC | ATTTGGGGAGCAATGGAGGA |
| α‐SMA (1) | CCCCTGAAGAGCATCGGACA | TGGCGGGGACATTGAAGGT |
| α‐SMA (2) | GACGTACAACTGGTATTGTG | TCAGGATCTTCATGAGGTAG |
| TGF-β1 | TCAGACATTCGGGAAGCAGT | ACGCCAGGAATTGTTGCTAT |
| Kloth | TGGTTCGCCAACCCCATCCA | TGGGCCCGAAGGAAAAGGCA |
| Wnt-1 | ATTCTTCTTCTGGGGTGGGG | CCCCACCCCAGAAGAAGAAT |
| Snail-1 | TCCACAAGCACCAAGAGTCT | AGACTCTTGGTGCTTGTGGA |
| IL-1β | GAAATGCCACCTTTTGACAGTG | CTGGATGCTCTCATCAGGACA |
| IL-6 (1) | AGAAGGAGTGGCTAAGGACC | TGGTCCTTAGCCACTCCTTC |
| IL-6 (2) | CCAGTTGCCTTCTTGGGACT | GGTCTGTTGGGAGTGGTATCC |
| TNF-α | CAGGCGGTGCCTATGTCC | CGATCACCCCGAAGTTCAGTAG |
| Vimentin | CGGCTGCGAGAAATTGC | CCACTTTCCGTTCAAGGTCAAG |
| CTGF | GTGCCAGAACGCACACTG | CCCCGGTTACACTCCAAA |
| MPC1 | TAAAAACCTGGATCGGAACCAA | GCATTAGCTTCAGATTTACGGGT |
| GAPDH | AGGTCGGTGTGAACGGATTTG | GGGGTCGTTGATGGCAACA |
| β-Actin | TGTGGATCGGTGGCTCCATCCT | AAACGCAGCTCAGTAACAGTCCGC |
