## Supplementary material for "Artesunate Alleviate Kidney Fibrosis by Restoring Klotho Protein and Suppressing Wnt/β-Catenin Signalling Pathway": Suppl.Fig

**Supplementary Figures**

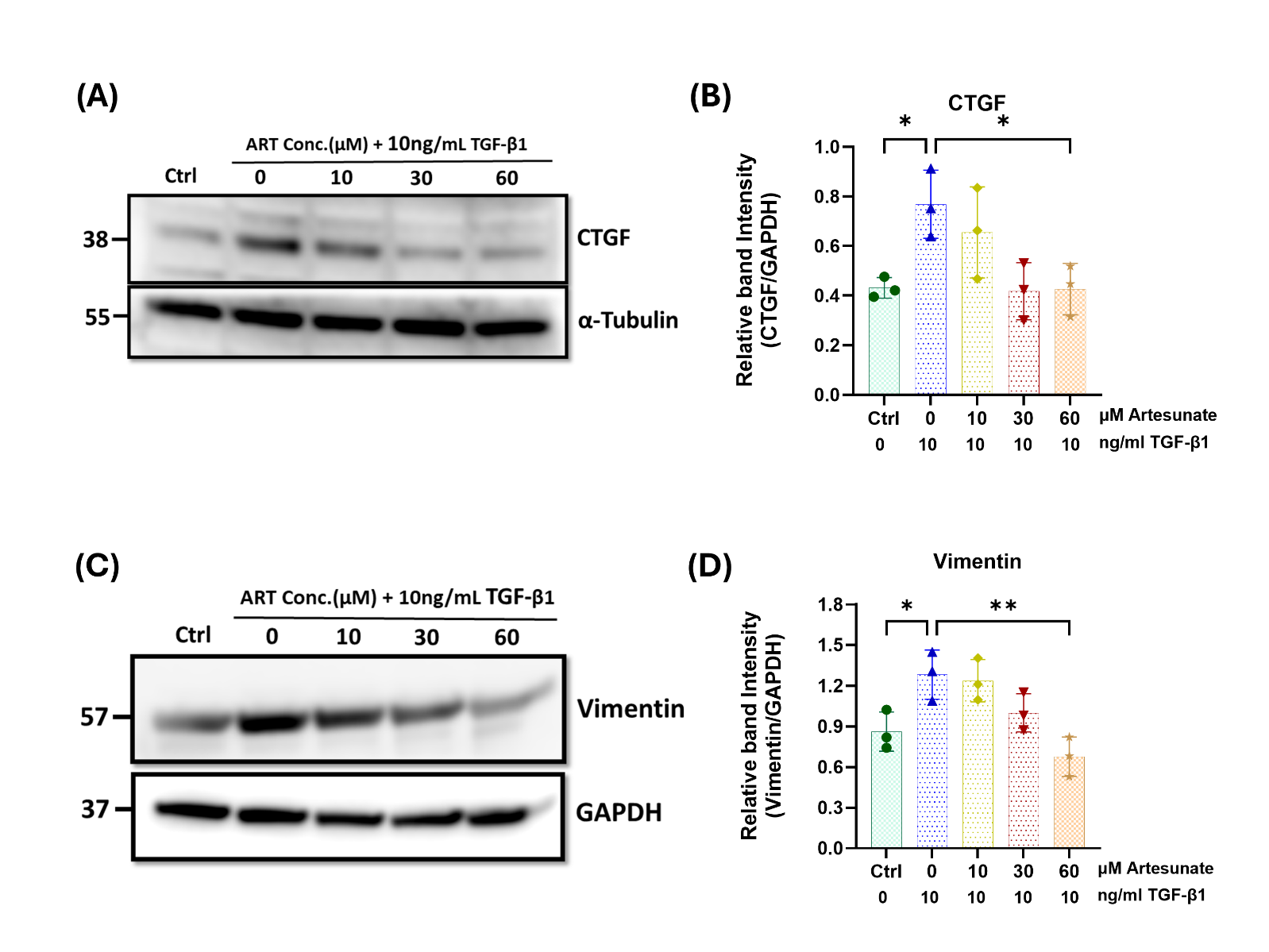

**Supplementary figure 1: Artesunate alleviated pro-fibrotic markers and reduced deposition of ECM and collagen in primary human kidney fibroblast (HKF) cells.** HKF cells were grown in DMEM supplemented with 10% FBS, 1% penicillin/streptomycin and the cells were maintained in a humidified atmosphere of 20% O2, 5% CO2 at 37 °C. The cells were starved with serum free medium for 24h and then stimulated with 10ng/ml TGF-β1, before treatment with different concertation of artesunate ranged from 0-60µM for 48h. The expression level of CTGF **(A,B)** and vimentin **(C,D)** proteins in respond to the different concentrations of artesunate were evaluated using western blotting technique and the blots were quantified by using ImageJ software and normalised to the GAPDH (n=3). Bars represent the means ± SEM from three independent experiments. One-way ANOVA was used for statistical analysis. *P<0.05, **p<0.01, ***p<0.001.

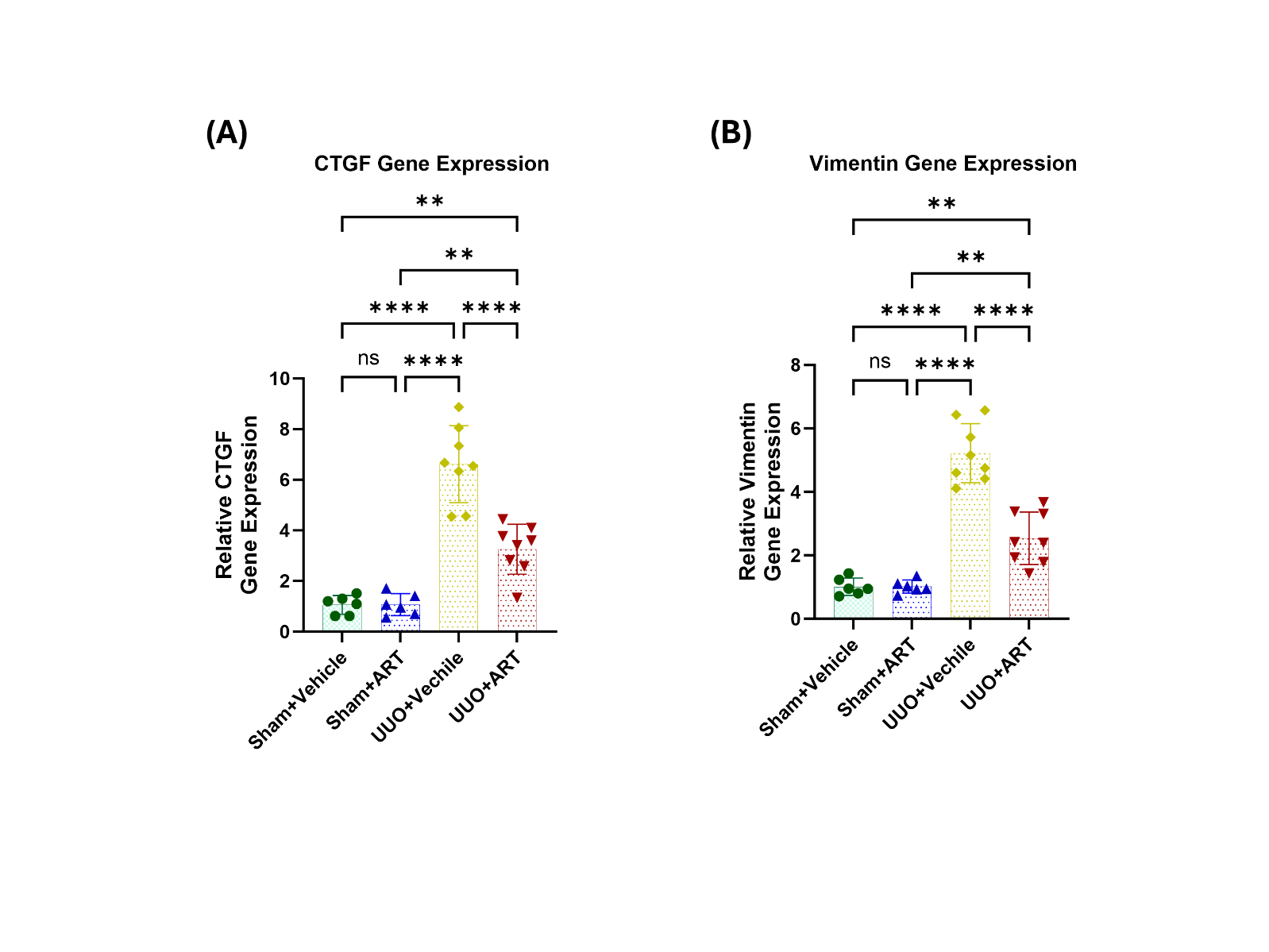

**Supplementary figure 2: Artesunate treatment attenuated of pro-fibrotic markers gene expression. (a)** and **(B)** Representative of relative gene expression of CTGF and vimentin from kidney of animals subjected to UUO, treated with artesunate or vehicle as indicated verse control group. For all graphs, error bars represent the means ± SEM of data from 6–8 animals per group. One-way ANOVA was used for statistical analysis. *P<0.05, **p<0.01, ***p<0.001.

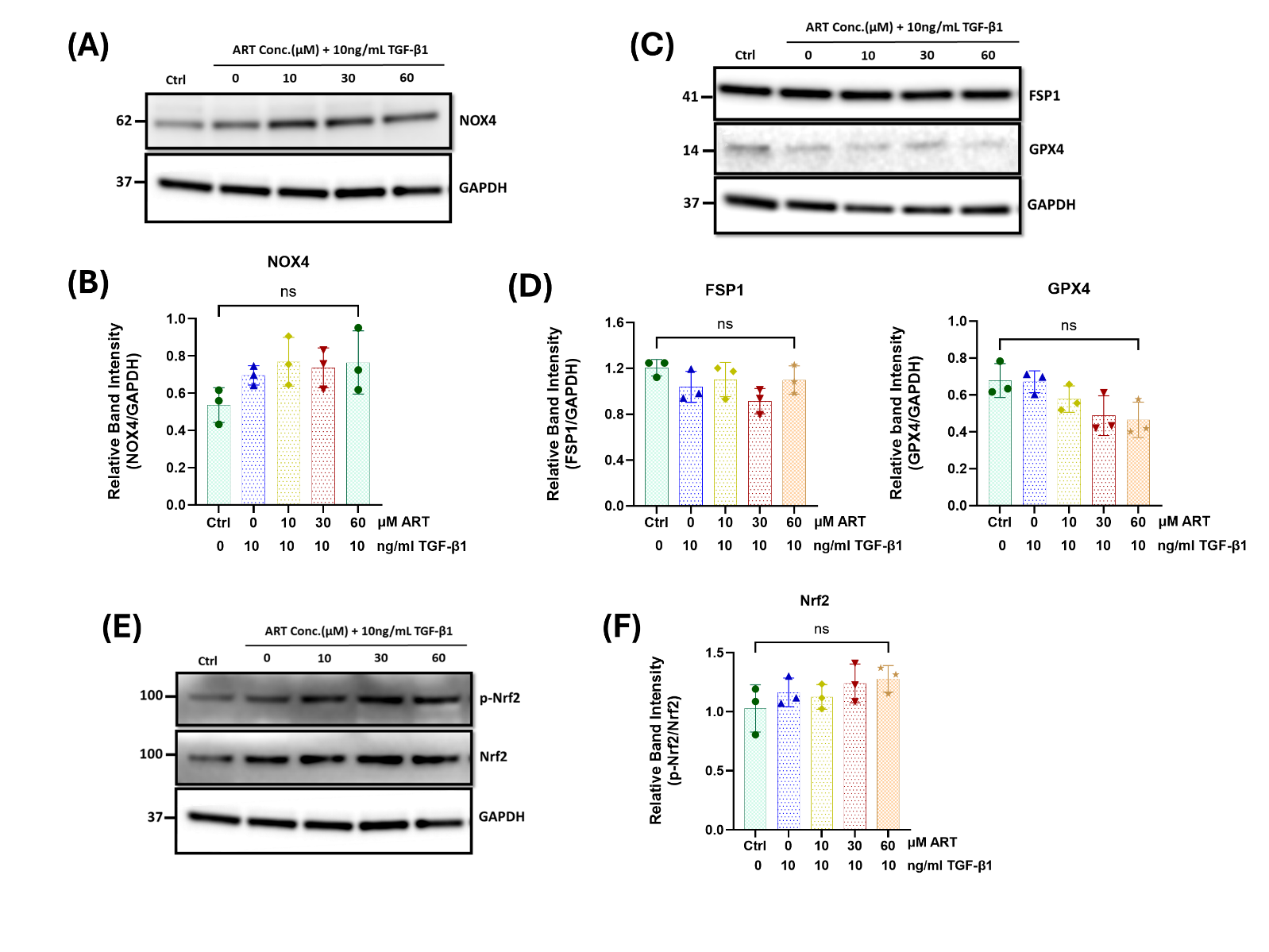

**Supplementary figure 3. Artesunate treatment did not increase oxidative stress and did not induce ferroptosis in human renal proximal tubule cells (HK2).** Serum starved human renal proximal tubule cells were stimulated with 10ng/ml TGF-β1 before treatment with different concertation of artesunate ranged from 0-60μM for 48h. The cells were then lysed with RIPA buffer containing protease and phosphatase inhibitor cocktails. Protein quantity was measured using BCA method and equal protein quantity was used for western blotting analysis. The expression level of NOX4 **(A)**, FPS1 and GPX4 **(B),** and p-Nrf2 and Nrf2 **(C)** in respond to the different concentrations of artesunate were evaluated using western blotting technique and the blots were quantified buy using ImageJ software and p-Nrf2 normalised to the total Nrf2 protein, and all other proteins normalised to the GAPDH (n=3). Bars represent the means ± SEM from three independent experiments. One-way ANOVA was used for statistical analysis. *P<0.05, **p<0.01, ***p<0.001.
